# Novel compound inhibitors of HIV-1_NL4-3_ Vpu

**DOI:** 10.1101/2021.07.30.454560

**Authors:** Carolyn A. Robinson, Terri D. Lyddon, Hwi Min Gil, David T. Evans, Yury V. Kuzmichev, Jonathan Richard, Andrés Finzi, Sarah Welbourn, Lynn Rasmussen, N. Miranda Nebane, Vandana V. Gupta, Sam Ananthan, Zhaohui Cai, Elizabeth R. Wonderlich, Corinne E. Augelli-Szafran, Robert Bostwick, Roger G. Ptak, Susan M. Schader, Marc C. Johnson

## Abstract

HIV-1 Vpu targets the host cell proteins CD4 and BST-2/Tetherin for degradation, ultimately resulting in enhanced virus spread and host immune evasion. The discovery and characterization of small molecules that antagonize Vpu would further elucidate the contribution of Vpu to pathogenesis and lay the foundation for the study of a new class of novel HIV-1 therapeutics. To identify novel compounds that block Vpu activity, we developed a cell-based ‘gain of function’ assay that produces a positive signal in response to Vpu inhibition. To develop this assay, we took advantage of the viral glycoprotein, GaLV Env. In the presence of Vpu, GaLV Env is not incorporated into viral particles, resulting in non-infectious virions. Vpu inhibition restores infectious particle production. Using this assay, a high throughput screen of >650,000 compounds was performed to identify inhibitors that block the biological activity of Vpu. From this screen, we identified several positive hits but focused on two compounds from one structural family, SRI-41897 and SRI-42371. It was conceivable that the compounds inhibited the formation of infectious virions by targeting host cell proteins instead of Vpu directly, so we developed independent counter-screens for off target interactions of the compounds and found no off target interactions. Additionally, these compounds block Vpu-mediated modulation of CD4, BST-2/Tetherin and antibody dependent cell-mediated toxicity (ADCC). Unfortunately, both SRI-41897 and SRI-42371 were shown to be specific to the N-terminal region of NL4-3 Vpu and did not function against other, more clinically relevant, strains of Vpu.

## INTRODUCTION

Since the beginning of the HIV epidemic, more than 75 million people have been infected with HIV and about 33 million people have died as a result. While there have been advances in treating HIV, approximately 1 million people per year die from complications resulting from HIV infection, and about 38 million people are still living with HIV/AIDS (American Foundation for Aids Research, 2020). Importantly, HIV can remain infectious at low levels ‘hidden’ in a host, despite modern advances in combinational Antiretroviral Therapies (ART). Furthermore, current therapies require life-long, daily treatment, which is expensive and unattainable for many of those living with HIV. Importantly, if drug treatment ceases, viral loads increase rapidly, resulting in a restoration of pre-treatment viral load levels that will continue HIV spread and eventually lead towards the development of AIDS (1). As eradication of HIV is unlikely without a cure, research efforts to further the understanding and control HIV are of utmost importance.

A potential target of future therapies is Vpu. Vpu is an HIV-1 accessory protein involved in viral assembly but not absolutely required for viral replication *in vitro*. Vpu is between 81-86 amino acids long and has been well characterized in its role of downmodulating CD4 to prevent superinfection and antagonizing BST-2/Tetherin to facilitate viral release (2–4). Additionally, Vpu has been reported to have other cellular targets including natural killer, T and B cell antigen (NTB-A) (5), CD155 (also known as polio virus receptor (PVR)) (6), CCR7 (7), CD62L and SNAT1 (8), although these targets are less characterized.

Vpu contains a Transmembrane domain (TMD) at its N-terminus and a cytoplasmic tail domain (CTD) at its C-terminus. Much of the activity of Vpu has been shown to be dependent on the phosphorylation of two serine residues in the cytoplasmic tail (S52/56) that are phosphorylated by casein kinase-2 (CK-2) and used to interact with a cellular ubiquitin ligase (9, 10). Specifically, Vpu depends on the phosphorylation of two serine residues for its ability to interact with a member of the Skp1-cullin1-F-box complex (SCF) containing the specific F-box proteins βTrCP-1 and βTrCP-2 and connect to the proteasome degradation pathway (10). Our lab has previously shown that in a cell line containing a CRISPR/Cas9 double knockout of the βTRCP proteins (1 and 2), Vpu activity against CD4 was abolished, and anti-BST-2/Tetherin activity was reduced significantly (11). This suggests that Vpu has at least one additional mechanism used for counteracting BST-2/Tetherin. The S52/56 mutations also no longer affected Vpu activity against BST-2/Tetherin in the knockout cell line. This is consistent with previous findings that Vpu and BST-2/Tetherin interact using their transmembrane domains (12, 13).

The transmembrane domain of Vpu has been shown to be required for the antagonism of BST-2/Tetherin, specifically residues A14/18 (14). Additionally, several poorly characterized targets of Vpu have been shown to be targeted by the transmembrane domain (A14/18 dependent) rather than the cytoplasmic domain, including NTB-A, PVR, and CD62L (5, 6, 15, 16). Targets of the transmembrane domain are generally thought to be downmodulated through mis-trafficking rather than degradation; however, this has yet to be fully elucidated (13). Kueck, et. Al., 2015, suggested that Vpu interacts with AP1/AP2 Catherin adapters to facilitate the endosomal degradation of BST-2/Tetherin in a manner that is also dependent on the S52/56 phosphoserines.

In addition to its well characterized role in targeting cellular proteins, Vpu has been shown to function as an ion channel for monovalent cations such as sodium and potassium (17, 18). While this function appears to be conserved, it is poorly understood and has yet to be linked to pathogenicity of HIV-1.

While most HIV-1 infected patients do not produce enough neutralizing antibodies to control the infection, the immune system can mediate the elimination of HIV-1 infected cells through antibody dependent cellular cytotoxicity (ADCC). During an ADCC response, antibodies bind to the surface of infected cells using a foreign antigen and recruit effector cells, including natural killer (NK) cells, that lyse the infected cell, eliminating virus-producing cells. In the case of HIV-1, the antigens on the surface of an infected cell are the envelope glycoproteins (Env). Several groups have shown that Vpu is an important player in protecting HIV-1 infected cells from lysis by ADCC (19, 20). CD4 downregulation prevents Env-CD4 interaction at the surface of infected cells, thus limiting the exposure of CD4-induced Env epitopes targeted by ADCC-mediating, non-neutralizing antibodies naturally present in the plasma from HIV-1-infected individuals (21, 22). Downregulation of BST-2/Tetherin, which otherwise traps viral particles at the surface of infected cells, contributes to limit the overall amount of Env on the surface of infected cells and to a lower ADCC response (22–24).

The impact of Vpu on viral spread suggests considerable therapeutic potential for a Vpu-specific inhibitor (25, 26). There have been several attempts to specifically target Vpu; however, many of the proposed compounds target the ion-channel activity of Vpu and have been shown to have low selectivity index or non-Vpu-specific ion-channel targets (18, 27, 28). Notably, the compound BIT225 is currently in Phase II of the Australian New Zealand clinical trials and has been shown to increase plasma derived activated CD4^+^ and CD8^+^ T cells and NK cells if coupled with ART, when compared with patients being treated with ART alone (29). However, it is important to note that BIT225 is not Vpu specific (28, 30) and does not appear to affect Vpu-mediated antagonism of BST-2/Tetherin (31). There have been additional efforts to target the ubiquitin ligase machinery upstream of Vpu, such as with the compound MLN4924, but this compound does not rescue ADCC activity at tolerated levels in cells (32).

Finally, our lab has previously demonstrated that Vpu downmodulates Gibbon Ape Leukemia Virus Envelope (GaLV Env) through targeting of the cytoplasmic tail region using the same SCF/βTrCP ubiquitin ligase (11, 33, 34). Using this observation, we developed a novel assay for a gain-of-function high throughput screen for inhibitors of Vpu. Here, we identify a family of novel inhibitors specific to NL4-3 HIV-1Vpu. Two of the identified compounds, SRI-41897 and SRI-42371, were examined further and found to effectively block Vpu downmodulation of GaLV Env, BST-2/Tetherin, CD4, and rescue ADCC killing of infected cells.

## MATERIALS & METHODS

### Plasmids

NL4-3 derived HIV-1-CMV-GFP was provided by Vineet Kewal Rammani (National Cancer Institute (NCI) – Frederick) and previously described (35). This proviral vector lacks the accessory genes *vif, vpr, nef*, and *env* and contains a CMV promoter driven GFP in the place of *nef*. Both Vpu positive and Vpu negative constructs were used and described previously (33). The infectious molecular clone NL4-3 was obtained through the NIH HIV Reagent Program, Division of AIDS, NIAID, NIH, contributed by M.A. Martin (36).

HIV-1 Gag-mRFP constructs were created in a two-step cloning process. First, the Nhe1-Xho1 region of NL4-3 derived HIV-1-CMV-GFP constructs (+/- Vpu) containing GFP were replaced with a Blasticidin Resistance gene gblock fragment ordered from Integrated DNA Technologies (IDT). Secondly, the BamH1-Xho1 region of a previously described Gag-mRFP construct gifted by Akira Ono (University of Michigan) was replaced with the BamH1-Xho1 region of the new constructs where Gag-mRFP replaces Pol (37).

The MLV/GaLV chimera construct used in infectivity assays was described previously (11, 33, 34). The APOBEC3G-GFP was gifted by Daniel Salamango and described previously (38). Plasmids for creation of stable cell lines were created by insertion of CDC25α-GFP or MLV/GaLV into the retroviral transfer vector pQCXIH (Clonetech, Mountain View, CA, USA). The SHIV_KU-1vMC33_ strain of Vpu was gifted from Edward Stephens (University of Kansas), and the CH77 strain Vpu was inserted using a GBlock (IDT). Both Vpu strains were cloned into the NL4-3 derived HIV-1-CMV-GFP plasmid described above. The proline insertion mutant was created using PCR mutagenesis of NL4-3 Vpu and subsequent infusion into the NL4-3 derived HIV-1-CMV-GFP plasmid described above.

### Compounds

Compound SRI-40244 (T6143675, CAS # 1090371-79-1) and Compound SRI-41897 (Z285895332, CAS # 1208824-23-0) were purchased from Enamine. Compound SRI-42371 was purchased from Ambinter (Amb11288305 CAS # 2185475-57-2). Compounds were resuspended at 40 mM (stock concentration) in DMSO and stored at 4°C.

### Cell Lines

The base HEK293FT (293FT) cell line used for stable expression was originally obtained from Invitrogen (Carlsbad, CA, USA). For the GaLV inhibition assay, cell lines stably expressing GaLV/MLV chimera construct were transduced with NL4-3 derived HIV-1-CMV-GFP virus pseudotyped with VSV-G.

Counter-screen stable cell lines (AOBEC3G-GFP and CDC25Cα-GFP) were cloned into a pQCXIH retroviral transfer vector packaged and transduced into HEK293FT cells using VSV-G gylocprotein, then selected with hygromycin. In both cases, a single cell isolate was selected and propagated for use.

The 293T mCAT-1 cell line expressing the ecotropic F-MLV Env receptor was provided by Walther Mothes.

TZM-GFP cells were gifted by Massimo Pizzato and previously described (39).

All 293FT based cell lines and TZM-GFP cells were maintained in Dulbecco’s Modified Eagle Medium (DMEM) supplemented with 7.5% fetal bovine serum, 2mM L-glutamine, 1mM sodium pyruvate, and 10mM nonessential amino acids.

The cell lines for the ADCC assay were obtained and cared for as previously described (40). The NK cell line (KHYG-1 cells) was obtained from the Japan Health Sciences Foundation and transduced with the V158 variant of human CD16. The target cells (CEM.NKR-CCR5) were obtained from the AIDS Reagent Program and were modified as described by transducing with a vector carrying an SIV LTR driven luciferase reporter gene.

Primary human peripheral blood mononuclear cells (PBMCs) and CD4^+^ T cells were isolated, activated, and cultured as previously described (41). Briefly, PBMCs were obtained by leukapheresis and CD4^+^ T lymphocytes were purified from resting PBMCs by negative selection using immunomagnetic beads per the instructions of the manufacturer (StemCell Technologies, Vancouver, BC, Canada) and were activated with phytohemagglutinin-L (10 μg/mL) for 48 h and then maintained in RPMI 1640 complete medium supplemented with recombinant interleukin-2 (rIL-2) (100 U/mL). VSV-G-pseudotyped HIV-1 NL4-3 virus was produced and titrated. Viruses were then used to infect activated primary CD4 T cells from healthy HIV-1-negative donors by spin infection at 800 × *g* for 1 h in 96-well plates at 25°C. **Ethics Statement.** Written informed consent was obtained from all study participants and research adhered to the ethical guidelines of CRCHUM and was reviewed and approved by the CRCHUM institutional review board (ethics committee, approval number CE16.164-CA). Research adhered to the standards indicated by the Declaration of Helsinki. All participants were adults and provided informed written consent prior to enrolment in accordance with Institutional Review Board approval.

### GaLV Assay/screen/flow

Infectivity (GaLV) Assay. Cell lines stably expressing GaLV/MLV chimera construct were transduced with NL4-3 derived HIV-1-CMV-GFP virus pseudotyped with VSV-G, and treated with compound, DMSO (Sigma-Aldrich Cat# D4540), or MLN4924 (Calbiochem Cat# 5.05477.0001) for 24 hours at 37°C. Viral media was collected and frozen at −80°C for between 2 and 24 hours. Once thawed, virus was spun at 3000 RCF for 5 minutes to clear any debris. Viral supernatant (NL4-3 derived HIV-1-CMV-GFP pseudotyped with GaLV ENV/MLV chimera) was then used to transduce mCAT-1 cells for 48 hours at 37°C and collected for flow cytometry. GFP positive mCAT-1 Cells (indicating a successful transduction) were measured.

### High Throughput Screen

#### Compound collection

Southern Research maintains a collection of 759,059 unique, non-proprietary compounds assembled from various commercial vendors (Enzo, Selleck, ChemBridge, Enamine, Life Sciences) for screening targets in HTS. Eight molecular properties were calculated for this collection using Accelrys Pipeline Pilot application. Analysis showed that 89.7 % of compounds have molecular properties matching all eight criteria for lead-like molecules to serve as starting points for a drug discovery effort (Molecular Weight <= 500; Heteroatom count <=10; Number Rotatable Bonds <= 8; Number Aromatic Rings <= 4; A Log P < = 6; Molecular Polar Surface Area <= 200; H-bond acceptors < 10; H-bond donors < 5). Compound collection: Southern Research maintains a collection of 759,059 unique, non-proprietary compounds assembled from various commercial vendors (Enzo, Selleck, ChemBridge, Enamine, Life Sciences) for screening targets in HTS. Eight molecular properties were calculated for this collection using Accelrys Pipeline Pilot application. Analysis showed that 89.7 % of compounds have molecular properties matching all eight criteria for lead-like molecules to serve as starting points for a drug discovery effort (Molecular Weight <= 500; Heteroatom count <=10; Number Rotatable Bonds <= 8; Number Aromatic Rings <= 4; A Log P < = 6; Molecular Polar Surface Area <= 200; H-bond acceptors < 10; H-bond donors < 5). Within this chemical space, the collection is diverse containing: 1. 278,767 non-overlapping Murcko scaffolds with an average cluster size 2 to 3, 2. 9,228 individual ring systems with unique substitution pattern (average frequency 218) and, 3. 16,008 contiguous ring systems with unique substitution patterns (average cluster frequency 125). A subset of 674,336 compounds from this collection were tested in HTS format at a single concentration of 10 μg/mL or 30 μM depending on the compound library source.

#### Assay Method

Library compounds were diluted in assay medium (DMEM with 10% FBS, 1% PSG, 1% HEPES) to prepare a 3.5x concentrated dosing solution (35 μg/mL or 105 μM) and added to 384-well black clear bottom plates (Corning; Cat # 3764BC) in 10 μL (1/3.5 final well volume). Twenty-five μL of Producer cells (HEK 293 FT cells stably expressing GaLV Env and pNL4-3 ΔEnv-GFP) at 800,000 Cells/mL was added to the plates for a final count of 20,000 Cells/well, a final compound concentration of 10 μg/mL or 30 μM, and a final DMSO concentration of 0.5%. Assay medium alone (at 0.5% DMSO) served as the negative control (columns 1 and 2 of each plate) and 0.5 μM MLN4924 (Activebiochem; Cat # MLN4924) (at 0.5% DMSO) as the positive control (columns 23 and 24 of each plate). The plates were incubated at 37°C/5% CO_2_ for 24 h in a humidified atmosphere. Ten μL of supernatant was transferred from each plate into a new 384-well black clear bottom plate, and 20 μL of Acceptor cells (293FT cells stably expressing mCAT-1 and the MLV Env receptor) at 300,000 Cells/mL added to the plates now containing 10 μL supernatant for a final count of 6,000 Cells/well. The plates were returned to 37°C/5% CO_2_ for 72 h and GFP signal imaged on a Mirrorball plate reader (BMG Labtech). The fluorescence signals of the test wells were normalized to percent activation relative to the average of the positive control wells on each plate by the formula:

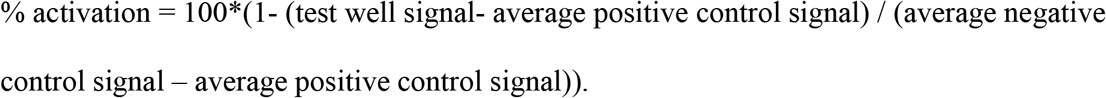

Potential hits from this primary HTS were cherry-picked and tested at ten concentrations in 2-fold serial dilution over a concentration range of 50 – 0.1 μM or μg/mL, in both the main assay outlined above as well as in a counter screen assay.

#### HTS Counter Screen

Compounds were diluted in assay medium (DMEM with 10% FBS, 1% PSG, 1% HEPES) to prepare a 3.5x concentrated dosing solution and added to 384-well black clear bottom plates (Corning; Cat # 3764BC) in 10 μL (1/3.5 final well volume). Twenty-five μL of Acceptor cells (293FT cells stably expressing mCAT-1 and the MLV Env receptor) was added to the plates for a final count of 6,000 Cells/well, a final compound concentration of 50 – 0.1 μM or 50 – 0.1 μg/mL, and a final DMSO concentration of 0.5%. Assay medium alone (at 0.5% DMSO) and 0.5 μM MLN4924 (at 0.5% DMSO) served as controls. The plates were incubated at 37°C/5% CO_2_ for 24 h in a humidified atmosphere, after which GFP signal was imaged on a Mirrorball plate reader (BMG Labtech).

### Cellular Target Counter Screens

293FT cells stably expressing CDC25α-GFP or APOBEC3G-GFP (described above) were treated with 40 μM compound, DMSO, or MLN4924 for 24 hours at 37°C and collected for flow cytometry. Mean Fluorescence Intensity (MFI) was measured.

### CD4 and Tetherin Surface Labeling in PBMCs

PBMCs from HIV-negative donors were CD8-depleted (Invitrogen Cat# 11147D) and activated with 0.5μg/mL purified PHA (Remel Cat# R30852801) in complete media (RPMI-1640 with glutamax (Gibco Cat# 61870-036) containing 10% FBS (Peak Serum Cat# PS-FB1), 1% HEPES (Gibco Cat# 15630-080), and 1% Pen/Strep (Gibco Cat# 15140-122)) supplemented with 20 IU/mL rhIL-2 (R&D Systems Cat# 202-IL) for 72±4 hours at 37°C and 5% CO_2_. PHA was removed and washed out with complete media. 2.5×10^6^ P were resuspended in either 5 mL of VSV-g pseudotyped HIV-1_NL4-3_Δ_Env-eGFP_Δ_Vpu_, HIV-1_NL4-3_Δ_Env-eGFP_Δ_Nef_Δ_vpu_ or HIV-1 _NL4-3_Δ_Env-eGFP_Δ_Nef_ viral supernatants or infection media control (DMEM with glutamax (Gibco Cat# 10566-016) with 10% FBS, and 1% MEM Non-Essential Amino Acids (Gibco Cat# 11140-050)), aliquoted into 24 well plates, and spinoculated at room temperature for 2 hours at 2000 rpm with no brake applied. Following spinoculation, viral supernatant was removed and replaced with 1mL complete media with 20 IU/mL rhIL-2. After 24±4 hours, compounds were added to each well at the final concentration of 25 μM. Following additional incubation for 24±4 hours at 37°C and 5% CO_2_, supernatant was removed, and cells were treated with an antibody cocktail containing 1:400 Live/Dead stain (InVitrogen Cat# L34965), 1:100 CD4 stain (BioLegend Cat# 300530, clone RPA-T4), and 1:100 BST-2/Tetherin stain (BioLegend Cat# 348415, clone RS38E) in 3% FBS in PBS (Gibco Cat# 14190144) in the dark at 4°C for 30 minutes. Cells were washed 3x with 3% FBS in PBS, fixed in 2% PFA (Affymetrix Cat#199431LT) in PBS, and analyzed using flow cytometry on Stratedigm S1000Exi (Stratedigm, Inc).

### CD4 Surface Labeling in TZM-GFP

TZM-GFP cells were transduced with VSV-G pseudotyped packaged HIV-1 constructs containing either *gag* and *env* and lacking *vif, vpr, pol* and *vpu* or containing *gag* and *env* and lacking *vif, vpr, pol* and *vpu*. Both constructs contain mCHERRY in place of the Pol gene. Day 4 post infection, media was changed, and cells were treated with 40μM SRI-41897, SRI-42371, 1μM MLN4924, or 0.1% DMSO for 24 hours at 37°C and 5% CO_2_. For surface staining, cells were lifted using TrypLE Express and transferred to round bottom 2 mL tubes in 1mL of PBS. Cells were pelleted at 300 RCF for 3 minutes and supernatant was removed. Cells were resuspended in a blocking solution containing 5% goat serum in PBS at 4°C for 30 minutes. Cells were pelleted at 300 RCF for 3 minutes and blocking solution was removed. Staining was done in a 1% goat serum and PBS solution containing 1:100 APC conjugate CD4 Stain (Life Technologies REF# MHCD0405) at 4°C for 1 hour in the dark. Cells were pelleted at 300 RCF for 3 minutes, and stain was removed. Cells were washed 3x in 1mL of PBS. Following washes, cells were fixed in 1% PFA in PBS. PFA solution was removed after centrifugation for 3 minutes at 300 RCF, and cells were resuspended in 400μL of PBS and analyzed on n Accuri C6 flow cytometer.

### Flow Cytometry

Cells were washed with PBS and lifted using Tryple Express (Gibco). Cells were removed from the plate and moved to 1.5 mL microcentrifuge tubes containing PBS and Paraformaldehide (PFA) at a final concentration of 4% PFA for 10 minutes at room temperature. Following, cells were spun down at 800 RCF for 5 minutes and washed 2x with PBS before the final resuspension in PBS and analyzed using an Accuri C6 flow cytometer.

### ADME

Standard kinetic solubility assay was performed at pH 7.5 (simulated intestinal fluid) using an LC-MS method to quantify the samples and Estradiol (haloperidol) as a control standard. Solubility testing was done in triplicate. The distribution coefficient (LogD) was calculated using Microsoft Excel following quantification of samples using a LC-MS method. The assay was run at pH 7.5 (PBS buffer) and used Verapamil as a control standard. Testing was done in triplicate. For microsomal stability testing, compounds were incubated with liver microsomes (from mouse or human) in the presence of NADPH. Compound concentrations were then measured via LC-MS/MS at 0, 5, 10, 20, and 45 min. The half-life or intrinsic clearance of each compound was then calculated. Diclofenac was used as a positive control.

### 2G12 Surface Labeling

Surface Labeling 2G12 was previously described (42). Infected primary CD4 T cells were treated with compound 24 hours post infection. 48 hours post infection, cells were incubated for 20 min at 37°C, with 5 μg/mL 2G12 (AB002; Polymun). Cells were then washed once with PBS and incubated with 1 ug/ml anti-human (Alexa Fluor 647; Invitrogen) secondary Abs and the viability dye AquaVivid (Thermo Fisher Scientific) for 15 min in PBS. Cells were washed again with PBS and fixed in a 2% PBS-formaldehyde solution. Infected cells were stained intracellularly for HIV-1 p24, using a Cytofix/Cytoperm fixation/permeabilization kit (BD Biosciences, Mississauga, ON, Canada) and fluorescent anti-p24 MAb (phycoerythrin [PE]-conjugated anti-p24, clone KC57; Beckman Coulter/Immunotech). The percentage of infected cells (p24^+^) was determined by gating the living cell population on the basis of viability dye staining (Aquavivid; Thermo Fisher Scientific). Samples were acquired on an LSR II cytometer (BD Biosciences), and data analysis was performed using FlowJo vX.0.7 (Tree Star, Ashland, OR, USA).

### ADCC

The ADCC assay was previously described (40). Target cells were infected 4 days prior to each assay with either WT NL4-3 HIV-1 or NL4-3 HIV-1 ΔVpu. CEM.NKR-CCR5 cells were infected by spinoculation for 2 hours at 1,200 RCF. After spinoculation, virus was removed, and target cells were cultured in R10 medium. Immediately before the assembly of ADCC assays, infected target cells were washed three times in R10 medium. 24 hours after infection, compounds were added to the infected target cells at a concentration of 25 μM for each independently. 48 hr after infection, target cells were washed then incubated with the NK cell line KHYG-1 at a ratio of 10:1 of 150,000 effector cells to 15,000 target cells in round-bottom, tissue culture-treated polystyrene 96-well plates. Assays were performed in R10 culture medium containing 10 U IL-2 per mL, with no CsA. Each plate contained NK effector cells and uninfected target cells in the absence of antibody (0% relative light units (RLU)), and NK cells and infected targets in the absence of antibody (100% RLU). Serial, 4-fold, triplicate dilutions of plasma or monoclonal antibody were added. Each compound was once again used to treat the cell mixture at 25 μM. Once targets, effectors, and serially diluted antibody were combined, assay plates were incubated for 8 h at 37°C and 5% CO2. After an 8h incubation, a 150-μl volume of cells was resuspended and mixed by pipette with 50 μl of the luciferase substrate reagent BriteLite Plus (Perkin Elmer) in white 96-well plates. Luciferase activity was read approximately 2 min later using a Wallac Victor3 plate reader (Perkin Elmer). 50% ADCC titers were estimated as previously described for virus neutralization assays (41). The 50% intercept was calculated using the adjacent %RLU values above and below 50% RLU.

## RESULTS

### High Throughput Screen for Vpu Inhibitors

Our lab previously showed that Vpu prevents infectious particle production in HIV-1 particles pseudotyped with a GaLV Env chimera consisting of Murine Leukemia Virus (MLV) Envelope containing a GaLV Env c-terminal-domain (CTD) (henceforth referred to as GaLV Env) that is targeted by Vpu for downmodulation (33). Using this observation, we developed a gain-of-function high throughput screen (HTS) for Vpu inhibitors (Figure 1A). This screen uses HEK 293 FT cells stably expressing GaLV Env and pNL4-3 ΔEnv-GFP to produce virus followed by transduction into 293FT cells stably expressing mCAT-1, the MLV Env receptor, as target cells (34). The cells produce HIV-1 viral particles pseudotyped with GaLV Env that successfully transduce mCAT-1 expressing cells in the absence of Vpu and produce few to no viral particles if the NL4-3 provirus contains a functioning Vpu (Figure 1B). The compound MLN4924 is a Neddylation inhibitor that effectively blocks the function of the cullin-Ring group of ubiquitin ligases and therefore blocks the cellular machinery that Vpu depends on for much of its activity (10, 11, 32, 43). In the absence of Vpu or in the presence of an inhibitor blocking Vpu, such as MLN4924, infectious particle production is rescued (Figure 1B).

**Figure 1:**
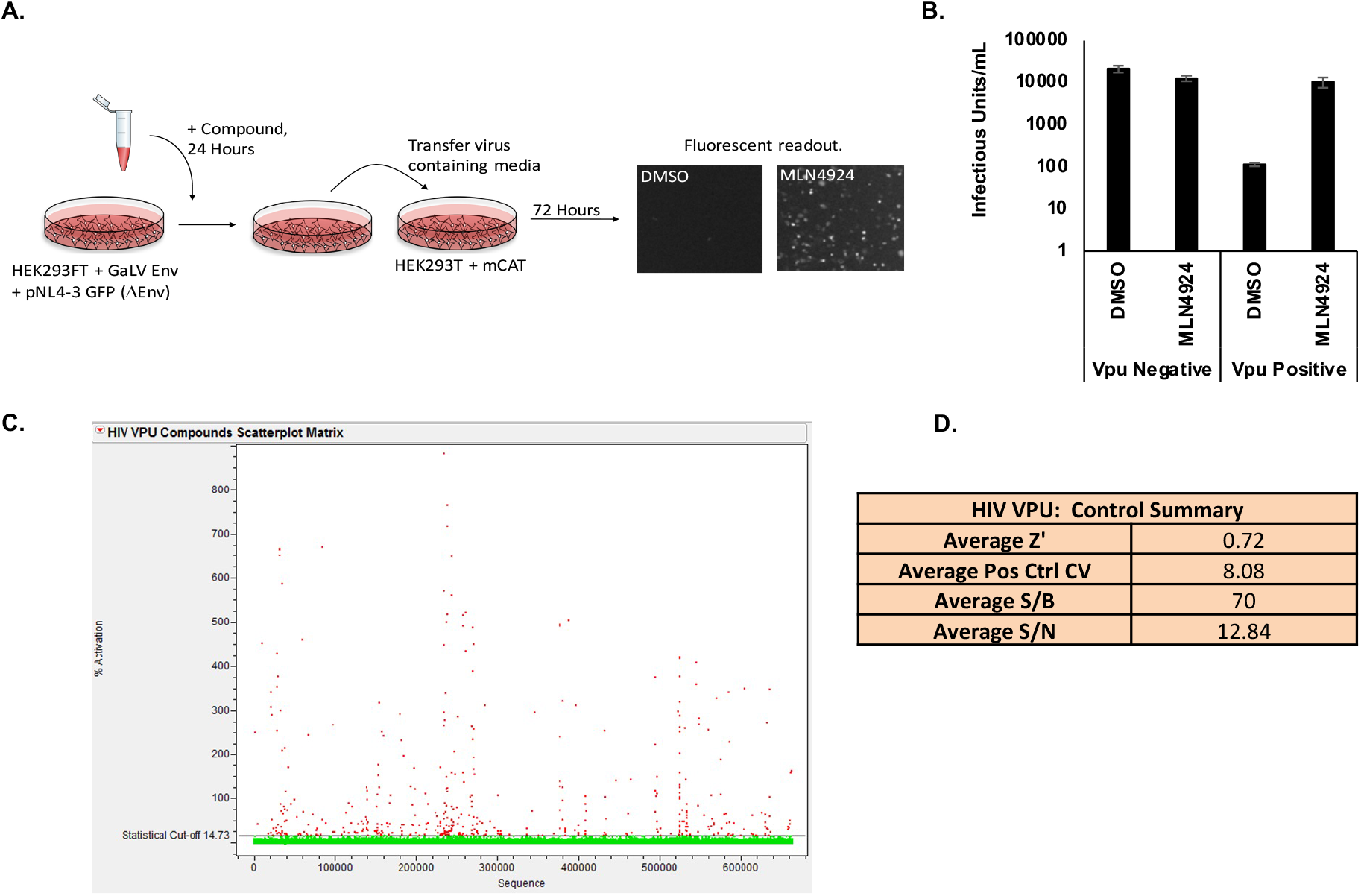
High Throughput Screen for Vpu Inhibitors. **A)** Schematic of GaLV Infectivity Assay experimental design. Cells stably expressing GaLV/MLV Env were transduced with a HIV-1 provirus. Cells were treated with compound for 24 hours, and viral media was used to infect cells expressing MLV receptor. Vpu blockage results in higher infectivity. **B)** GaLV Infectivity Assay. Loss of infectious particle production is Vpu dependent and is rescued by MLN4924. Data is represented on a logarithmic scale. N=3, and error bars represent standard deviation. **C)** Scatter plot of HTS data. The threshold for selecting active compounds (red symbols) is 14.73 % (line) as determined by the mean + 3xSD of all the data. **D)** Statistical values showing HTS assay performance determined from high (positive) and low (negative) control values on each plate. Z’ = 1 - 3(StdDevHiControl +StdDevLowControl) /(AvgHiControl - AvgLowControl); % Pos Cont CV = 100 * StdDevHiControl / AvgHiControl; S/B (Signal/Background) = Avg HiControl / AvgLowControl; S/N (Signal/Noise)=(AvgHiControl - AvgLowControl) / sqrt(StdDevHiControl^2^ + StdDevLowControl^2^).

This assay was used in the HTS to screen 674,336 commercially available compounds for Vpu inhibition (Figure 1C and D). The average Z’-value calculated from the 32 positive and 32 negative control wells on each plate was 0.72, with a range from 0.51 to 0.85, for the 2145 assay plates run in the screen. A Z’-value > 0.5 indicates that the assay performance was adequate to detect active compounds when tested once (44). Statistical analysis (mean+3xSD of all test compound data) identified 14.73% inhibition as the cutoff between inactive and active compounds. Based on this cutoff, a total of 461 compounds were identified as hits resulting in an overall hit rate of 0.07% for the primary screen. Of the 461 hit compounds identified, 202 were confirmed as active following retest at ten concentrations in the GFP reporter assay used in HTS. These compounds were also evaluated for concentration-response in counter screen assays measuring compound autofluorescence or non-Vpu mediated activation of GFP expression. Based on these results, 177 compounds were excluded as causing autofluorescence or having off-target effects. The remaining 25 compounds were selected for further testing in secondary assays of which 21 were available for purchase as fresh powders.

### HTS Reveals Potent Vpu Inhibitors

One compound, SRI-41897, inhibited Vpu more potently than others. Interestingly, SRI-41897 had a very similar structure to another of the 21 HTS hits, SRI-40244, highlighting the potential for the structural family as Vpu inhibitors (Figure 2A, B). While multiple occurrences of similarly structured compounds is interesting, SRI-40244 had very low activity in the GaLV inhibition assay and was not perused further (data not shown). Interested in this structural class of compounds, we screened more than 80 additional commercially available analogs for SRI-41897 and found SRI-42371 that inhibited Vpu more potently in the GaLV Inhibition assay than the original compound (Figure 2C,D). These two compounds became the focus of our study. The EC_50_ of both compounds is in the μM range, 4.4 and 7.4 μM for SRI -41897 and SRI-42371, respectfully, as measured in the GaLV inhibition assay (Figure 2E,F).

**Figure 2:**
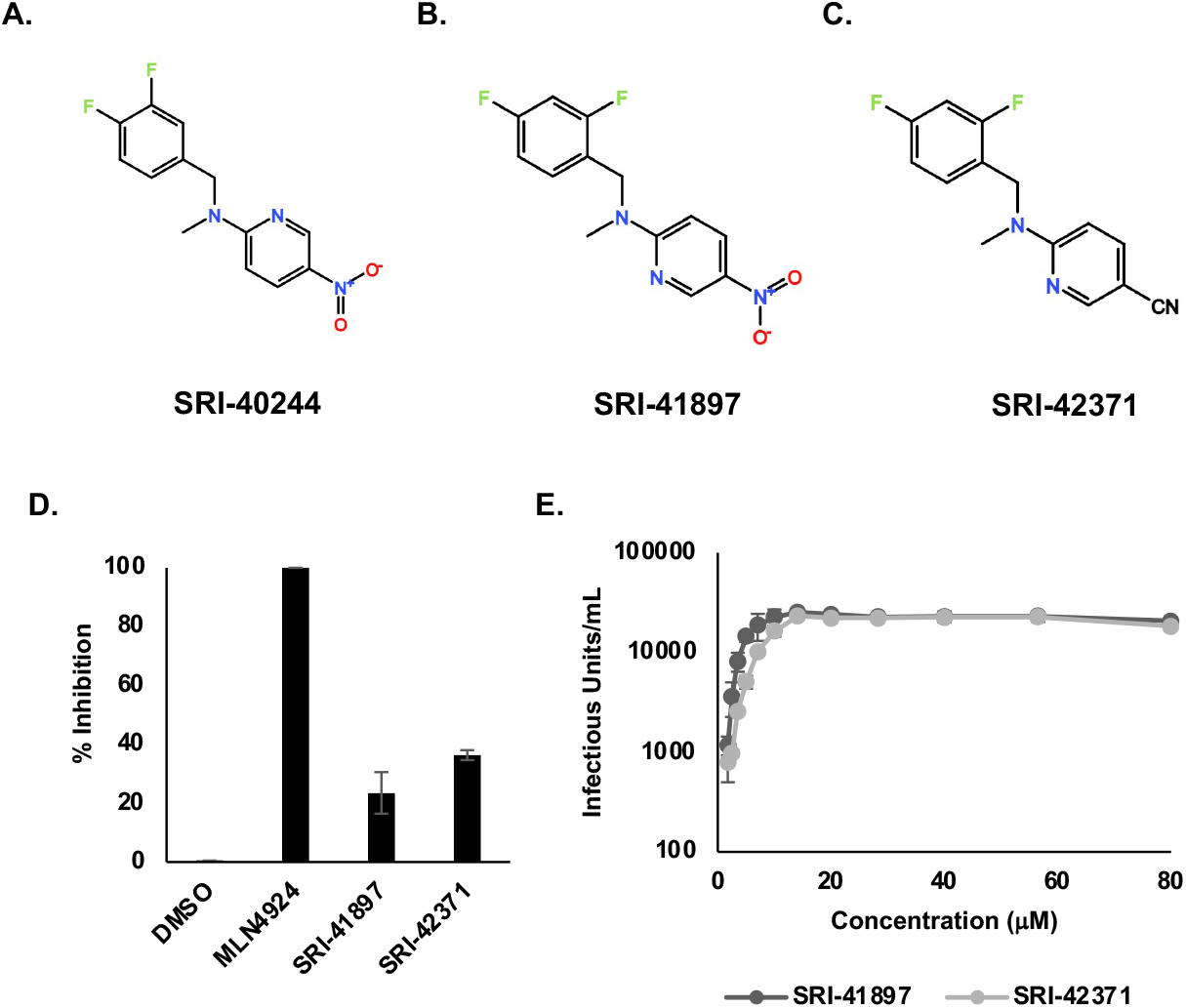
HTS reveals potent Vpu Inhibitors. **A)** Structures of SRI-40244, **B)** SRI-41897, **C)** SRI-42371 drawn on TouchMol Structure drawing program available online. **D)** GaLV infectivity assay rescue shown normalized to MLN4924. Compounds added at 40μM in ‘semistable’ transduced cells as described in Figure 1A. Error bars reflect standard deviation, N=4. **E)** EC50 was determined using the GaLV infectivity assay (Figure 1A) for compound concentrations ranging from 0.88 to 80 μM. Error bars reflect standard deviation. N=3. The EC50 was calculated using the AAT Bioquest EC50 calculator available online. The calculated EC50 for SRI-41897 and SRI-42371 were 4.4 μM and 7.4 μM, respectively.

### Vpu-Independent Counter Screens

While the HTS is Vpu dependent, compounds that block the cellular machinery that Vpu depends on will also result in a positive hit. To ensure that the compounds are specific for Vpu, we developed a series of counter screens for the upstream cellular machinery maintaining MLN4924 as a positive control (Figure 3A).

**Figure 3:**
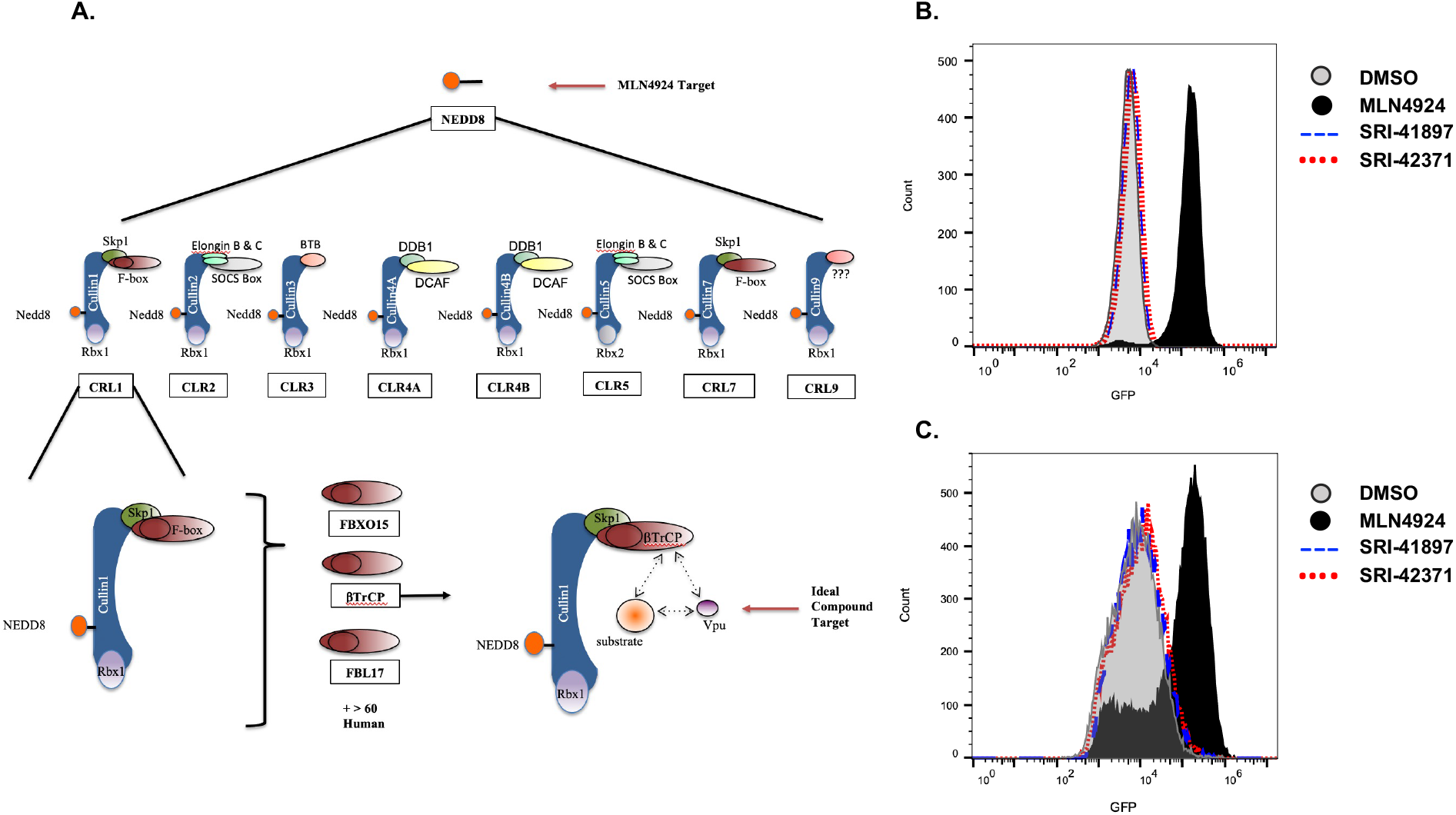
Vpu-Independent Counter Screens. **A)** Schematic of potential upstream targets of Vpu inhibitors. **B)** Cells expressing GFP-tagged APOBEC3G and HIV-1 Vif were treated were treated with either 40mM of SRI-41897, SRI42371 or one of the controls, DMSO or MLN4924 for 24 hours and collected for flow cytometry. Mean fluorescence intensity (MFI) is shown in representative images. **C)** Cells expressing GFP-tagged CDC25α were treated with either 40mM of SRI-41897, SRI42371 or one of the controls, DMSO or MLN4924 for 24 hours and collected for flow cytometry. Mean fluorescence intensity (MFI) is shown in representative images.

The first counter screen ensures that the compounds do not have any activity for the broader cullin family of ubiquitin ligases. In this assay, 293FT cells that are stably expressing both GFP tagged APOBEC3G and the HIV-1 accessory protein, Vif, were treated with compounds or MLN4924, and MFI was recorded via flow cytometry (Figure 3B). Vif has previously been shown to downmodulate the cellular protein APOBEC3G using cullin5 machinery (45–48), so a blockage of cullin5 results in an increase of APOBEC3G expression and therefore an increase in GFP as shown by MFI. Addition of MLN4924 showed a clear increase in MFI of GFP, whereas neither SRI-41897 nor SRI-42371 displayed an increase in fluorescence, indicating that they do not block cullin5 or the larger cullin ubiquitin ligase family. The second counter screen ensures that the compounds do not have any activity against the F-box protein, βTrCP 1 or 2. In this assay, 293FT cells stably expressing GFP tagged CDC25α were treated with compounds, DMSO, or MLN4924 as a positive control, and MFI was recorded via flow cytometry (Figure 3C). βTrCP has been previously shown to downmodulate the cellular protein CDC25α; therefore, a blockage of βTrCP results in an increase of CDC25α expression and an increase in GFP as shown by MFI (49, 50). Addition of MLN4924 resulted in a clear increase in fluorescence; however, neither SRI-41897 nor SRI-42371 displayed an increase in fluorescence, indicating that they do not block βTrCP function.

### Compounds Rescue CD4 and BST-2/Tetherin Expression

To explore the impact of compounds on surface expression of CD4, TZM-GFP cells were transduced with a Gag-mRFP construct containing a non-functional Pol protein and a functional Env in the presence or absence of Vpu (Figure 4A,B). CD4 levels of Vpu positive cells were only 6% of Vpu negative cells (Figure 4A). After treatment with MLN4924 or compounds, CD4 surface expression was rescued by 47% or 25% on infected TZM-GFP cells, for SRI-41897 or SRI-42371, respectively, in comparison to the negative control, DMSO (Figure 4B). While the compounds had no effect on CD4 surface expression in the absence of Vpu, it is interesting that MLN4924 reduced CD4 surface expression in these cells. This is likely why the rescue of CD4 surface expression with MLN4924 in the presence of Vpu was not significant.

**Figure 4:**
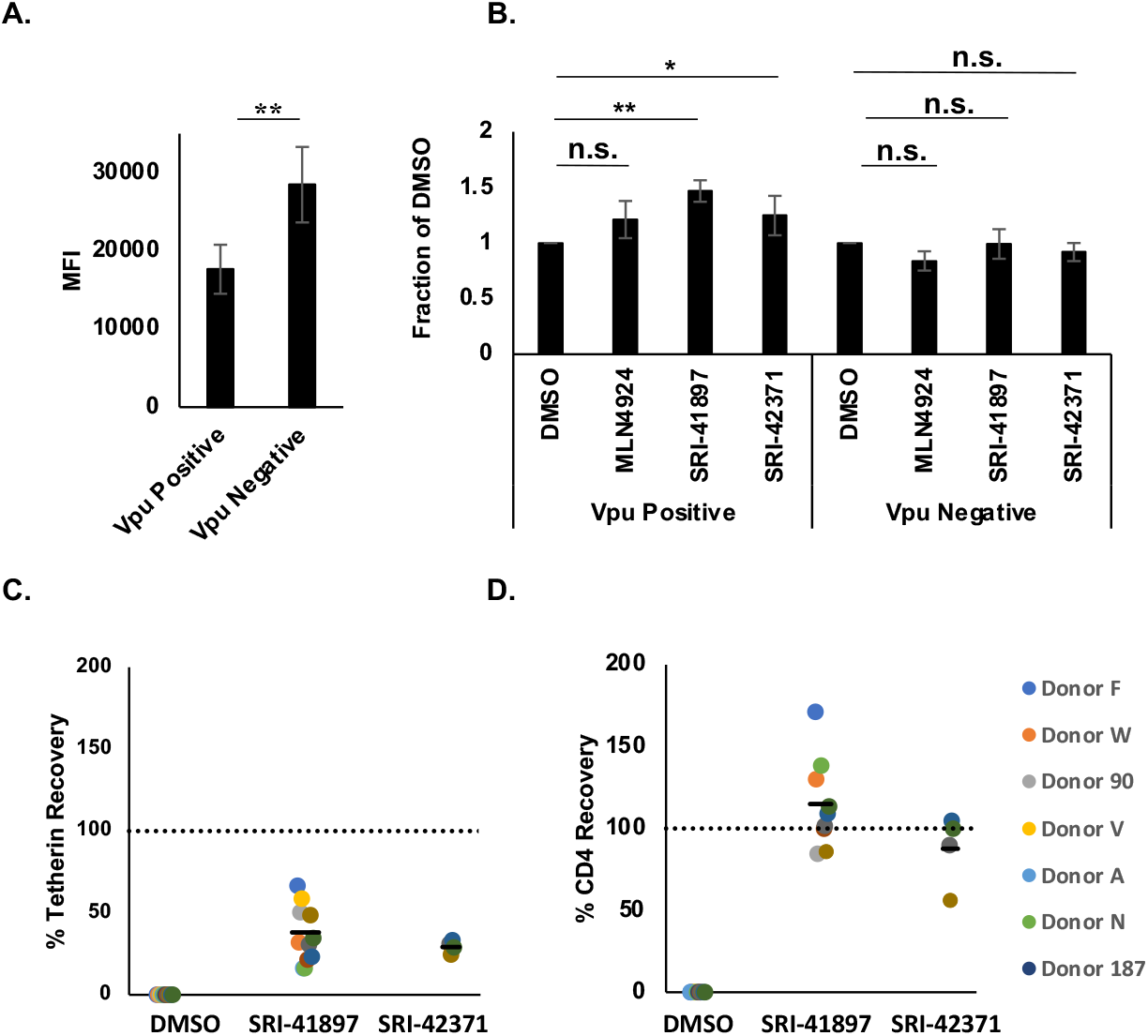
Compounds Rescue CD4 and Tetherin Expression. **A)** Surface Expression of CD4 in the presence and absence of Vpu. N=4. TZM-GFP cells were transduced with VSV-G pseudotyped packaged HIV-1 constructs containing mCHERRY. Day 4 post infection, media was changed, and cells were treated with 1μM MLN4924 or 0.1% DMSO for 24 hours then cells were surface for CD4 and analyzed by flow cytometry. Error bars represent standard deviation. **B)** Surface Expression of CD4 in the presence and absence of Vpu after treatment with compounds. N=4. Cells were surface labeled as in Figure 4A, with the addition of 40μM of SRI-41897 or SRI-42371. Data is normalized to DMSO. Error bars represent standard deviation. **C)** Tetherin surface labeling, and **D)** CD4 surface labeling were done as follows. PBMCs from HIV-negative donors were CD8-depleted and activated. Cells were resuspended in either VSV-g pseudotyped HIV-1_NL4-3_Δ_Env-eGFP_Δ_Vpu_, HIV-1_NL4-3_Δ_Env-eGFP_Δ_Nef_Δ_Vpu_ or HIV-1 _NL4-3_Δ_Env-eGFP_Δ_Nef_ viral supernatants or control media. After 24±4 hours, compounds were added to each well at the final concentration of 25μM for 24 hours. Following, supernatant was removed, and cells were surface labeled with an antibody cocktail containing Live/Dead stain, CD4 stain, and BST-2/Tetherin stain. Data was analyzed using flow cytometry. Black horizontal bars represent the mean recovery.

To determine if SRI-41897 and SRI-42371 can rescue the expression of CD4 and Tetherin in patient derived peripheral blood mononuclear cells (PBMCs), cells were infected with HIV-1_NL4-3_, treated with compound for 24 hours, and surface labeled. Surface levels of CD4 and BST-2/Tetherin were measured via flow cytometry. Tetherin expression was restored an average of 38% and 29% and CD4 expression was restored an average of 115% and 88% when treated with 25 μM SRI-41897 and SRI-42371, respectively, in PHA-activated, CD8-depleted PBMCs (Figure 4 C, D).

### Compounds Rescue ADCC Response

Antibody Dependent Cellular Toxicity (ADCC) is a cellular response to infection that results in cell killing through lysis. In recent years, several groups have shown that Vpu protects HIV-1 infected cells from killing by ADCC, and that this effect is mostly due to downmodulation of CD4 and BST-2/Tetherin (reviewed in (19, 51, 52)). We wanted to test if the compounds could rescue ADCC as they rescued the expression of CD4 and partial expression of BST-2/Tetherin (Figure 4). Before testing ADCC, we examined Env expression using a conformationally-independent anti-HIV-1 Env antibody 2G12 expression (53, 54). Primary CD4^+^ T cells were infected with WT NL4-3 virus, treated with compounds at various concentrations, and probed with the anti-HIV-1 Env antibody, 2G12 (Figure 5A). We observed a dose dependent response in 2G12 binding, indicating that in the presence of compound there is more Env present on the surface of the cell. Next, target cells derived from CEM.NKR-CCR5_CD4+_ T cells expressing a Tat-inducible luciferase reporter gene were infected with either WT NL4-3 HIV-1 or NL4-3 HIV-1 ΔVpu, co-cultured with natural killer (NK) cells and reciprocal anti-HIV-1 Immune Globulin (HIVIG) dilutions (40) (Figure 5B-D). In this assay, an increase in ADCC killing is shown through a decrease in luciferase signal (decrease in actively infected cells). In the absence of compound, ADCC levels are significantly higher in NL4-3 ΔVpu infected cells when compared with NL4-3 WT infected cells (Figure 5B); however, introduction of compound brought ADCC levels against NL4-3 WT ADCC close to ΔVpu levels (Figure 5C,D), indicating that the presence of compounds rescue ADCC killing in NL4-3 WT infected cells.

**Figure 5:**
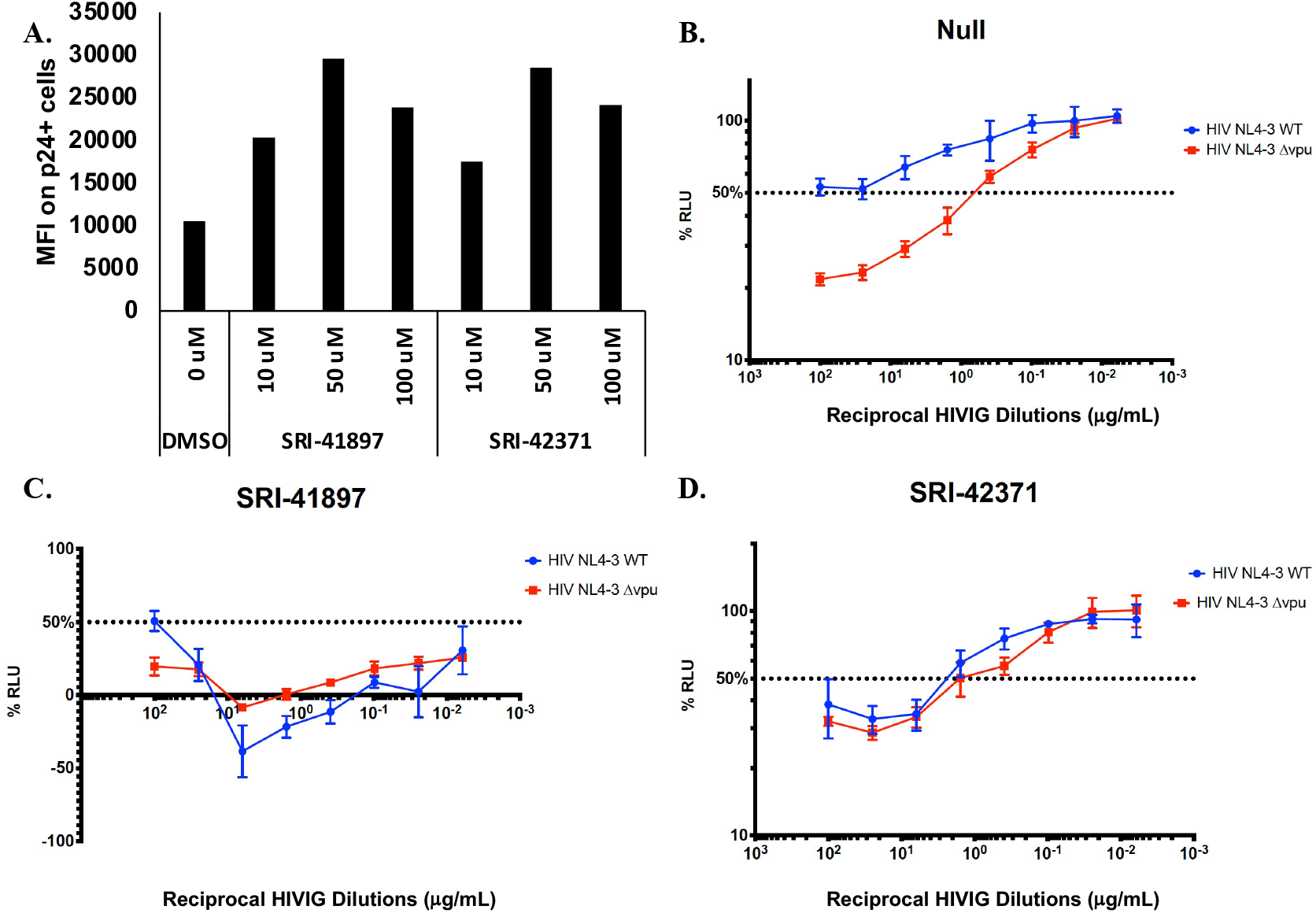
**A)** NL4-3 Infected primary CD4 T cells were treated with compound 24 hours post infection. After 48 hours post infection, cells were surface stained with 2G12 antibody. Additionally, infected cells were stained intracellularly for HIV-1 p24. Samples were analyzed using flow cytometry. **B-D)** Compounds were added to infected target cells at a concentration of 25 μM each, independently. After 48-hours post infection, target cells were incubated with NK cells. For controls, each plate contained NK effector cells and uninfected target cells in the absence of antibody (0% relative light units (RLU)), and NK cells and infected targets in the absence of antibody (100% RLU). Serial, 4-fold, triplicate dilutions of plasma or monoclonal antibody were added. Luciferase signal was detected in each well. The 50% intercept was calculated using the adjacent %RLU values above and below 50% RLU. Error bars represent standard deviation.

### Compounds are NL4-3 Strain Specific

To test the breadth of Vpu strains that are inhibited by SRI-41897 and SRI-42371, we tested inhibition against a wide variety of available Vpu strains and discovered a strain very similar to NL4-3 Vpu that was resistant to both compounds; the strain is from a simian-human immunodeficiency virus strain, SHIV_KU-1vMC33_, containing a Vpu originally from HIV-1 HXB2 (55, 56). SHIV_KU-1vMC33_ Vpu contains only 4 amino acid changes from NL4-3 Vpu (Figure 6A,C). Additionally, another, more clinically relevant strain of Vpu, CH77, was cloned into a NL4-3 backbone. While approximately 74% identical, there are several differences between CH77 and NL4-3 Vpu (Figure 6C). The NL4-3, CH77, and SHIV_KU-1vMC33_ strains of Vpu were examined in the GaLV inhibition assay (introduced in Figure 1a) in the presence and absence of compound or MLN4924 (Figure 6A). Interestingly, both compounds had no inhibition of CH77 Vpu or SHIV_KU-1vMC33_.

**Figure 6:**
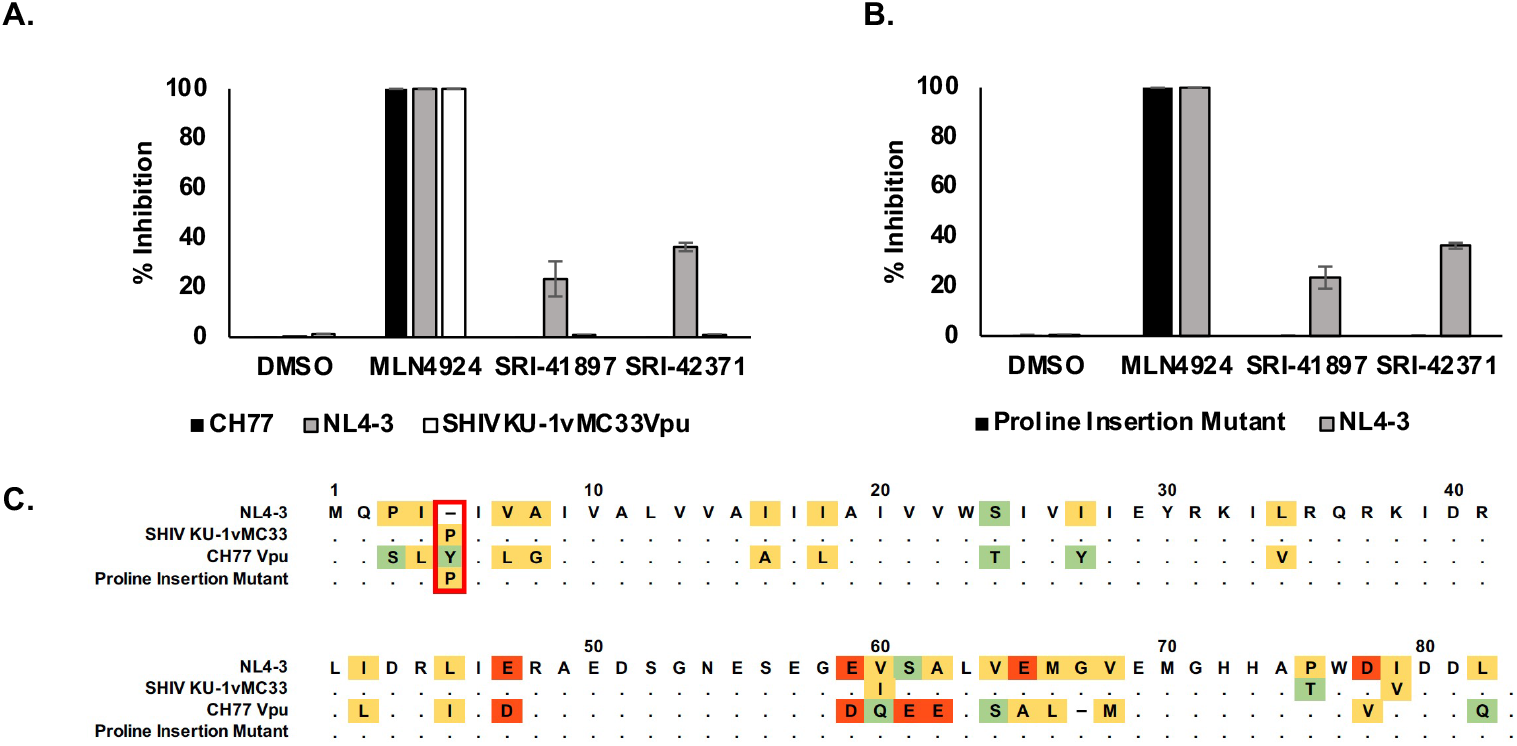
Compounds are NL4-3 Strain Specific. **A)** N=3. GaLV Infectivity Assay (Figure 1A) with NL4-3, CH77, and SHIV_KU-1vMC33_ Vpu strains cloned into a NL4-3 backbone and normalized to MLN4924. Error bars represent standard deviation. **B)** N=3. GaLV Infectivity Assay (Figure 1A) comparison of NL4-3 Vpu and proline insertion mutant normalized to MLN4924 positive control. Error bars represent standard deviation. **C)** Schematic of amino acid sequences of NL4-3, CH77, SHIV_KU-1vMC3_, and Proline Insertion Mutant Vpu strains. Highlighting reflects discrepancies in relation to NL4-3 Vpu.

The SHIV_KU-1vMC33_ Vpu and CH77 Vpu sequences differ in the same place from NL4-3 Vpu in only two places, AA5 and AA61 (Figure 6D). To examine the effect of this position on SRI-41897 and SRI-42371, a single proline, the same amino acid in position 5 of SHIV_KU-1vMC33_ was inserted into NL4-3 Vpu after amino acid 4 and tested alongside NL4-3 WT Vpu in the GaLV inhibition assay (Figure 6B). The single proline insertion was sufficient to completely abrogate the function of both compounds against Vpu. According to the Los Alamos database, less than 1% of Vpu contain single amino acid deletion between two side-by-side isoleucine residues at the N terminus like with NL4-3 Vpu. This suggests that the compounds would not likely be functional on most strains of Vpu.

## DISCUSSION

While still not fully elucidated, the inability to completely clear an HIV-1 infection could largely be due to accessory proteins, such as Vpu, that aid in immune system evasion through mechanisms such as downmodulation of CD4, BST-2/Tetherin, and inhibitory effects on ADCC. Currently, there are no approved therapies targeting HIV-1 Vpu.

Here we described two small molecules, SRI-41897 and SRI-42371, identified from a HTS, that inhibit HIV-1_NL4-3_ Vpu protein. The HTS took advantage of our previous finding that Vpu targets the cytoplasmic tail of GaLV Env using the same SCF/βTrCP ubiquitin ligase machinery it uses to downmodulate other targets (11, 33, 34). When HIV-1 particles are pseudotyped with GaLV Env in the presence of Vpu, almost no infectious viral particles are produced, but when Vpu is inhibited, infectious particle production is restored (Figure 1). This HTS has the advantage of being a gain-of-function assay reducing the chance of false-negative hits. Additionally, in a gain-of-function cellular screen, compounds must be able to cross the cell membrane to result in a positive hit, and highly toxic compounds will result in a negative signal.

The GaLV Inhibition HTS was used to screen 674,336 commercially available compounds and resulted in 21 positive hits. The 21 HTS hits included two compounds that were structurally similar, one of which, SRI-41897 had a strong signal. Over 80 commercially available analogs of SRI-41897 were screened in the GaLV inhibition assay, and a second compound, SRI-42371 more potently inhibited Vpu (Figure 2). While SRI-42371 inhibited Vpu more potently than SRI-41897 in the GaLV inhibition assay, the EC_50_ of SRI-42371 was higher. Furthermore, the nitro group present on SRI-41897 raises a concern of potential toxicity (Reviewed in (57)). Thus, SRI-42371 may be considered a better hit for optimization. With this in mind, SRI-41897 and SRI-42371 became the focus of our study moving forward.

Compounds of interest were counter screened in a Vpu-independent manner for auto fluorescence (data not shown) and inhibition of the SCF/βTrCP ubiquitin ligase machinery that Vpu depends on at two different levels. First, inhibition of the broader cullin family was tested using cells expressing GFP-tagged APOBEC3G in the presence of HIV-1 Vif (Figure 3b). Secondly, inhibition of cellular βTrCP 1 and 2 was tested in cells expressing GFP-tagged CDC25α (Figure 3c). Both SRI-41897 and SRI-42371 had a negative signal for auto-fluorescence, cullin-family inhibition, and βTrCP inhibition suggesting that they are specific to Vpu.

Next, we showed that SRI-41897 and SRI-42371 rescue CD4 and BST-2/Tetherin expression in PBMCs infected with HIV-1_NL4-3_ in a Vpu-dependent manner (Figure 4A, B). This result suggests that these two compounds hinder the functional activity of Vpu which is an important factor in evasion of the immune system during a HIV-1 infection. Furthermore, in recent years, Vpu has been shown to be an important factor in protecting infected cells from killing by ADCC (reviewed in (20)). Excitingly, we also showed that SRI-41897 and SRI-42371 increase the surface Env expression and killing of infected cells by ADCC (Figure 5).

It is important to note that part of the effect of Vpu on ADCC protection is linked to its ability to downregulate BST-2/Tetherin (22–24); however, the ability of Vpu to downregulate CD4^+^ also had significant effect on ADCC susceptibility (21, 58). This highlights the importance of a Vpu inhibitor that can rescue the expression of both BST-2/Tetherin and CD4 if it were to be used in a clinical setting.

The positive hit in the HTS, the negative hit in both counter screens, and CD4, BST-2/Tetherin, and ADCC rescue all suggest a dependence on Vpu. Unfortunately, we discovered that SRI-41897 and SRI-42371, specific to the NL4-3 strain of Vpu, are presumed to interact primarily with the N-terminus of Vpu around proline 4 and 5 (Figure 6). The addition of a proline after isoleucine 4 abrogated the effect of SRI-41897 and SRI-42371. While disappointing, the additional data showing NL4-3 specificity and dependance on a specific amino-acid sequence strongly suggest that both compounds are directly interacting with Vpu towards the N-terminus rather than any other cellular or HIV-1 targets. The finding that SRI-41897 and SRI-42371 interact with the N-terminus before or right at the transmembrane region is interesting because much of the activity of Vpu has been attributed to the cytoplasmic tail, specifically Serine 52/56. However, the transmembrane region downstream of where we hypothesize compounds are interacting, specifically A14/18, have been shown to be important for complete BST-2/Tetherin downmodulation, but this is not the case for CD4 downmodulation (11–14). An NMR structure of Vpu has been previously solved, and it is interesting to note that the group that changes in the transmembrane region of Vpu altered the angle of the cytoplasmic tail region (59). While further studies would be necessary, this is a possible explanation as to why a compound interacting at the N terminus of Vpu may disrupt functions associated with the cytoplasmic tail.

Even though SRI-41897 and SRI-42371 are NL4-3 specific, these two compounds have the potential to help further the understanding of Vpu and the effect its inhibition has on HIV-1 infection in both cell culture and animal models. Most importantly, we demonstrate here a solid proof-of-concept for high throughput screening for Vpu inhibitors. We were able to pull several small-molecule candidates from a large library of compounds, counter screen select candidate compounds, demonstrate functional inhibition, and locate important residues for these compounds to interact with Vpu. This gain-of-function assay may easily be adapted to screen inhibitors for other, more clinically relevant, strains of Vpu.

